# HuR-dependent expression of Wisp1 is necessary for TGF*β*-induced cardiac myofibroblast activity

**DOI:** 10.1101/2022.01.19.476976

**Authors:** Lisa C Green, Samuel Slone, Sarah R Anthony, Jeffrey Aube, Xiaoqing Wu, Liang Xu, Onur Kanisicak, Michael Tranter

**Author notes:** Correspondence: Michael Tranter, PhD, Division of Cardiovascular Health and Disease, Department of Internal Medicine, University of Cincinnati College of Medicine, 231 Albert Sabin Way, CVC 3928, Cincinnati, OH 45267.

## Abstract

Cardiac fibrosis is regulated by the activation and phenotypic switching of quiescent cardiac fibroblasts to active myofibroblasts, which have extracellular matrix (ECM) remodeling and contractile functions which play a central role in cardiac remodeling in response to injury. Here, we show that expression and activity of the RNA binding protein HuR is increased in cardiac fibroblasts upon transformation to an active myofibroblast. Pharmacological inhibition of HuR significantly blunts the TGF*β*-dependent increase in ECM remodeling genes, total collagen secretion, *in vitro* scratch closure, and collagen gel contraction in isolated primary cardiac fibroblasts, suggesting a suppression of TGF*β*-induced myofibroblast activation upon HuR inhibition. To delineate HuR-dependent mechanisms, we used photoactivatable ribonucleoside-enhanced crosslinking and immunoprecipitation (PAR-CLIP) to identify eleven mRNA transcripts that showed enriched HuR binding following TGF*β* treatment as well as significant co-expression correlation with HuR, *α*SMA, and periostin using single-cell RNA-sequencing from ischemic-zone isolated fibroblasts. Of these, Wnt1-inducible signaling pathway protein-1 (Wisp1; *Ccn4*), was the most significantly associated with HuR expression in fibroblasts. Accordingly, we found Wisp1 expression to be increased in cardiac fibroblasts isolated from the ischemic-zone of mouse hearts following ischemia/reperfusion, and confirmed Wisp1 expression to be HuR-dependent in isolated fibroblasts. Finally, addition of exogenous recombinant Wisp1 is able to partially rescue myofibroblast contractile function following HuR inhibition, demonstrating that HuR-dependent Wisp1 expression plays a functional role in HuR-dependent MF activity downstream of TGFβ. In conclusion, HuR activity is necessary for the functional activation of primary cardiac fibroblasts in response to TGF*β*, in part through post-transcriptional regulation of Wisp1.

**HIGHLIGHTS:**

- The RNA binding protein HuR is highly expressed in cardiac fibroblasts and its expression strongly correlates with markers of active myofibroblasts
- HuR inhibition reduces migration, contraction, and ECM production activity of cardiac fibroblasts
- Expression of the secreted matricellular protein, Wisp1, is increased in a HuR-dependent manner following TGFβ treatment
- Recombinant Wisp1 rescues myofibroblast contractile function following HuR inhibition
- HuR-dependent expression of Wisp1 is necessary for myofibroblast activation

## INTRODUCTION

Cardiac fibroblasts (CFs) are necessary for preserving physiological homeostasis in the myocardium, primarily through their central role in maintenance of the extracellular matrix (ECM). In the setting of cardiac stress or injury, quiescent CFs respond to chemical and mechanical stressors to undergo a phenotypic transformation to active myofibroblasts (MFs) [1,2]. Initial remodeling and biomechanical changes to the cardiac ECM may be a necessary and beneficial compensatory mechanism to allow the heart to maintain proper contractile function under this additional mechanical stress. However, chronic MF activation and excessive ECM remodeling is the driving force of cardiac fibrosis and is associated with increased progression to heart failure. Thus, a primary goal of the field is to identify pathways that can be targeted therapeutically to prevent excessive ECM remodeling resulting from chronic MF activity remains a goal.

We have previously shown that pharmacological inhibition of the RNA binding protein Human antigen R (HuR) reduced pathological cardiac remodeling, including a reduction in cardiac fibrosis, in a mouse model of cardiac pressure overload [3]. As part of this work, we demonstrated direct HuR-binding and post-transcriptional regulation of TGFβ expression in cardiac myocytes as a potential mechanism. TGFβ is the most common biochemical signaling initiator of MF activation and ECM remodeling activity [4,5]. In addition to our work showing TGFβ as a direct HuR target in cardiomyocytes, HuR has also been shown to promote TGFβ mRNA stability and expression in macrophages, a primary source of TGFβ following cardiac injury [6,7]. Interestingly, HuR has been suggested to facilitate a positive feedback loop of TGFβ signaling in primary rat cardiac fibroblasts as well [8]. This work also showed increased activity of HuR, as marked by increased cytoplasmic translocation, following TGFβ treatment. This increased activity was dependent on p38-MAPK, which we previously showed to also mediate HuR activation in hypertrophic myocytes [9]. More recently, HuR expression in macrophage-derived exosomes was shown to promote fibrosis in diabetic hearts, potentially through direct potentiation of fibroblast activity [10].

Together, the existing body of literature is suggestive of a functional role for HuR in mediating myofibroblast activation, but this has not yet been directly shown. Here, we utilized single cell RNA-sequencing from fibroblasts isolated from post-infarct mouse hearts to show that HuR expression is strongly correlated with existing myofibroblast marker genes. Inhibition of HuR in isolated primary CFs completely abrogates their ability to adopt a MF phenotype following TGFβ treatment, demonstrating that HuR directly mediates TGFβ-dependent MF activity. Photoactivatable ribonucleoside-enhanced crosslinking and immunoprecipitation (PAR-CLIP) was used to identify directly bound HuR target mRNAs in an unbiased, transcriptome-wide manner, and identified Wnt1-inducible signaling pathway protein-1 (Wisp1; *Ccn4*) as a novel TGFβ-dependent HuR target gene in fibroblasts. Wisp1 has been previously shown to increase in the myocardium following myocardial infarction and is suggested to mediate collagen production in isolated fibroblasts [11,12]. Finally, we show that restoring Wisp1 expression is sufficient to partially rescue TGFβ-dependent MF activity following HuR inhibition. In summary, these results demonstrate HuR-dependent expression of Wisp1 is both necessary and sufficient to mediate TGFβ-dependent MF activation.

## METHODS

### Isolation and culture of primary murine fibroblasts

Primary murine cardiac fibroblasts for culture experiments were isolated from adult wild-type C57BL/6 mice (8-12 weeks old). Fibroblasts were also isolated from the non-infarct and infarct zones of post-ischemic hearts of wild-type C57BL/6 mice (8-12 weeks old) following 30 minutes of ischemia and 14 days of reperfusion as previously described [13,14] to allow for mature scar/infarct formation. Following isolation, cardiac left ventricular tissue was minced in digestion buffer containing Collagenase IV (Worthington, LS004189) and Dispase II (Sigma, D4693) as described by Pinto et al [15]. The cell suspension was then incubated at 37°C for 60-90 minutes at 150 RPM on an orbital shaker with gentle disruption using a 10mL serological pipette every 15 minutes, followed by pre-plating for two hours in culture media (DMEM containing 10% FBS) to enrich for the fibroblast population. After two hours, the non-adherent cells were removed and the fibroblasts were given fresh culture media.

### Protein Isolation and Western blotting

Total protein was isolated from cells in RIPA buffer with 0.5 mM DTT, 0.2 mM sodium-orthovanadate, and protease inhibitor (Halt Protease and Phosphatase Inhibitor Cocktail; ThermoFisher). Protein extract (4 ug per lane) was separated on a 10% TGX Stain-Free polyacrylamide gel (Bio-Rad) and transferred to a nitrocellulose membrane, and loading was normalized to total protein as previously described [16,17]. Membranes were subsequently blocked for 1 h at room temperature using 5% Bovine Serum Albumin (BSA, MilliporeSigma, A7906) in 0.1% Tween 20, tris-buffered saline (T-TBS). Primary antibodies for Periostin (Novus Biologicals, NBP1-30042) and HuR (abcam, ab200342) were incubated overnight at 4°C at a dilution of 1:1000 in 4% BSA, and secondary antibody (anti-rabbit, BioRad 1721019), was incubated at a dilution of 1:5000 in 4% BSA for 1–2 hours at room temperature. Images were captured using chemiluminescent substrate (Thermo, 34580) on a ChemiDoc Imaging System and analyzed using ImageLab software (Bio-Rad).

### RNA isolation and qPCR

RNA was isolated using NucleoSpin RNA kits (Takara, 740955) and cDNA was synthesized using reverse transcription supermix for RT-qPCR (Biorad, 1708841). Quantitative PCR was performed using a BioRad CFX96 Real-Time System with iTaq Universal SYBR green supermix (Biorad, 1725121). Results were analyzed using the ***ΔΔ***Ct Method [18] using the following primers: 18S, F, 5’- AGTCCCTGCCCTTTGTACACA-3’; 18S, R, 5’- CCGAGGGCCTCACTAAACC-3’; *GAPDH*, F, 5’ ACCACAGTCCATGCCATCAC 3’, *GAPDH*, R, 5’ TCCACCACCCTGTTGCTGTA 3’, *postn*, F, 5’-GGTGGCGATGGTCACTTATT-3’, *postn*, R, 5’-GTCAGTGTGGTGGCTCTTAC-3’; *col1a1*, F, 5’-CTGGCGGTTCAGGTCCAA-3’, *col1a1*, R, 5’-GCCTCGGTGTCCCTTCATT-3’; *fn1*, F, 5’-GCTGGATAGATGGTGGACT-3’, *fn1*, R, 5’- AGCAGGTTCCCTCTGTTG-3’; *tgfb1*, F, 5’-CAATTCCTGGCGTTACCTTG-3’, *tgfb1*, R, 5’- CCCTGTATTCCGTCTCCTTG-3’; *ccn4*, F, 5’- GGTACGTCTCCCATGAGGTGGCTCCTGCCCT-3’, *ccn4*, R, 5’- CCGCCCCGACTCTAGAACTAATTCGCAATCTC TTCGAAGTCAGGGTAAGATTCCAA-3’.

### Immunohistochemistry

For *in vitro* IHC staining of primary fibroblasts, the media was aspirated and the cells were washed two times with PBS. The fibroblasts were then fixed in 4% paraformaldehyde (in PBS) for 15 min at room temperature, followed by 15 min incubations in 100% methanol, and then 70% ethanol. After rinsing in PBS, the cells were blocked in 6% Bovine Serum Albumin (BSA; Sigma Aldrich A3059) for at least an hour at room temperature or overnight at 4°C. Following blocking, cells were washed three times with PBS for 10 min at room temperature and incubated in primary antibody (anti-*α*SMA, Sigma, C6198) or anti-HuR, Abcam, ab28660) at 1:200 in 0.6% BSA for at least 2 hours at room temperature or overnight at 4°C. Cells were then incubated in AlexaFluor 488 anti-mouse secondary antibody (Abcam, ab150113) diluted 1:200 in 0.6% BSA for an hour at room temperature, washed 3 times with PBS, and incubated in Hoescht 33342 nuclear stain (1:3000 in PBS) for imaging of the nuclei.

### Scratch assay

Fibroblasts were plated on or before Passage 3 (P3) in a 96-well cell-culture dish. Upon reaching ~75% confluency (no more than 3 days), each well of fibroblasts was scratched using a P20 pipette tip to remove a strip of cells down the approximate center of the well. Following a single PBS wash, treatment media (e.g. HuR inhibitor and/or 10ng/mL TGFβ (Peprotech)) was added, and time-lapse images were obtained using the beacon function on a BioTek Cytation 5 multimodal imager. Scratch closure was quantified using the ImageJ “MRI_Wound_Healing_Tool” macro as described [19]. siRNA-mediated knockdown of HuR was achieved using 80 nM HuR targeting siRNA (Santa Cruz Biotechnology) transfected via Lipofectamine 3000 (ThermoFisher) as we have done previously [9] 48 hours prior to scratch assay.

### Collagen gel contraction

Collagen gels were made in a 24-well cell culture dish using a 1mg/mL collagen solution (Sigma, 08-115) and seeded with ~65,000 fibroblasts isolated from adult mouse hearts as described [20]. The seeded collagen gels were allowed to solidify at room temperature for 25 minutes and released from the sides and bottom of the well prior to treatment. Time-lapse imaging of gel contraction was performed and images were quantified using ImageJ.

### ECM decellularization

To allow for mature ECM deposition, primary fibroblasts were plated on or before P3 and grown in DMEM with 10% FBS and 1% penicillin/streptomycin for three days. The media was then changed to media containing vehicle or 10ng/mL TGFb1 (± KH-3) and cultured for an additional 3 days. Following a PBS rinse, decellularization and removal of soluble proteins was achieved by incubating in a solution of 0.25% Triton, 0.25% Na-Deoxycholate for 24 hours at 37°C as described [21].

### Picrosirius red staining

Cells were fixed in ice cold methanol overnight at −20°C, rinsed with PBS, and stained with Picrosirius Solution B (Polysciences, 24901B) for one hour at room temp. Wells were then rinsed twice with 0.1N HCl, and the picrosirius was eluted with 0.1M NaOH via gentle rocking at room temperature for one hour as described [22]. The resulting picrosirius elution was quantified using absorbance at 540nm on the BioTek Cytation 5.

### Mass spectrometry of decellularized ECM

Following decellularization as described above, ECM was washed three times with PBS and the remaining ECM matrix was solubilized with 2.5% (w/v) SDS in 50mM Tris pH=6.8 and mechanical scraping to ensure collection of all of the ECM as described [23]. The samples were then run 1.5cm into an Invitrogen 4-12% B-T gel using MOPS buffer with molecular weight marker lanes in between. The sections were excised, reduced with DTT, alkylated with IAA, and digested overnight with trypsin. The peptides were extracted, dried in a speed vac and resuspended in 7 ul of 0.1% formic acid, and 5.5 uL of each sample was analyzed by nanoLC-MS/MS (Orbitrap Eclipse). The combined data set of 9 injections were mapped against the mouse protein database using Proteome Discoverer v2.4 and the Sequest HT search algorithm (Thermo Scientific) with label-free quantitation (LFQ) parameters, but without any sample intensity normalization, by the UC Proteomics Core.

### PAR-CLIP

Fibroblasts were grown in 10cm dishes and treated with the photoreactive ribonucleoside 4-thiouridine (4SU; 0.1 mM) at both 20 hours and 4 hours to label long- and short-lived RNA transcripts, respectively, prior to RNA-protein cross-linking. Following a PBS rinse, cells were then crosslinked on ice at an energy setting of 200,000 uJ/cm^2^ using 368 nm wavelength bulbs. Protein G magnetic beads (Invitrogen) were washed five times in wash buffer (50mM Tris, 150mM NaCl, 1mM MgCl_2_, 0.05% NP40, pH=7.4) and loaded with 4 *μ*g of control (goat anti-rabbit; Invitrogen, 65-6120) or HuR antibody (Abcam, ab28660) in 0.02% Tween 20 in PBS for 30 minutes at RT, with gentle rocking [3]. Following cross-linking, cells were scraped into lysis buffer (100mM KCl, 5mM MgCl_2_, 10mM HEPES, 0.5% NP40, and protease inhibitor cocktail (ThermoFisher)), added to antibody-loaded Protein G beads, and incubated with 360° rotation at RT for 30 minutes. Following three washes using wash buffer to remove unbound RNA, bound RNAs were eluted via 30min incubation at 55°C in wash buffer supplemented with 0.5 *μ*g/*μ*L Proteinase K and 0.1% SDS and isolated using Qiazol/chloroform separation and NucleoSpin RNA columns (Takara). Luciferase control RNA is spiked in to each sample (5ng/sample) just prior to addition of Qiazol addition as an exogenous control for RNA yield and normalization of RNA-seq results, and only transcripts with a normalized transcript per million (TPM) value of ≥100 were considered HuR-bound targets.

### RNA-sequencing and bioinformatics analysis

RNA sequencing of paired end 100bp reads at a depth of ~30M reads/sample was performed by the Cincinnati Children’s Hospital Medical Center DNA Sequencing and Genotyping Core. Sequence read mapping and differential expression analysis was done using CLC Genomics Workbench v21.0.3 (Qiagen) as we have done previously [3,16,24]. The single cell RNA-seq dataset using isolated fibroblasts from post-MI hearts has been previously described and is publicly available (NCBI GEO Database accession GSE83337) [25]. Correlation analysis of gene co-expression within this dataset was done using transcriptome-wide Pearson correlation (statistical significance cutoff at P ≤ 0.05) modified from the WGCNA R-package (R v4.1.0) as previously described [24,26]

### Statistical analysis

All data was analyzed in a blinded fashion and presented as the mean with error bars representing standard error of the mean. Statistical comparisons between groups were done using Student’s t-tests or ANOVA, as appropriate, via GraphPad PRISM v9.2, with a p-value of less than 0.05 considered significant.

## RESULTS

### HuR expression and activation are increased upon myofibroblast activation

HuR protein expression in both CFs and MFs is much higher compared to non-stressed cardiomyocytes, where HuR appears largely absent from the blot at the same exposure time (Fig. 1A). Furthermore, analysis of protein expression shows that HuR expression appears to be increased in active MFs (treated with 10 ng/ml TGFβ for 24 hours) compared to quiescent CFs (Fig. 1A). Immuno-fluorescence (IF) staining shows increased cytoplasmic translocation of HuR (indicative of activation) in *α*SMA^+^ MFs 24 hours following TGFβ treatment (Fig. 1B).

**Figure 1.**
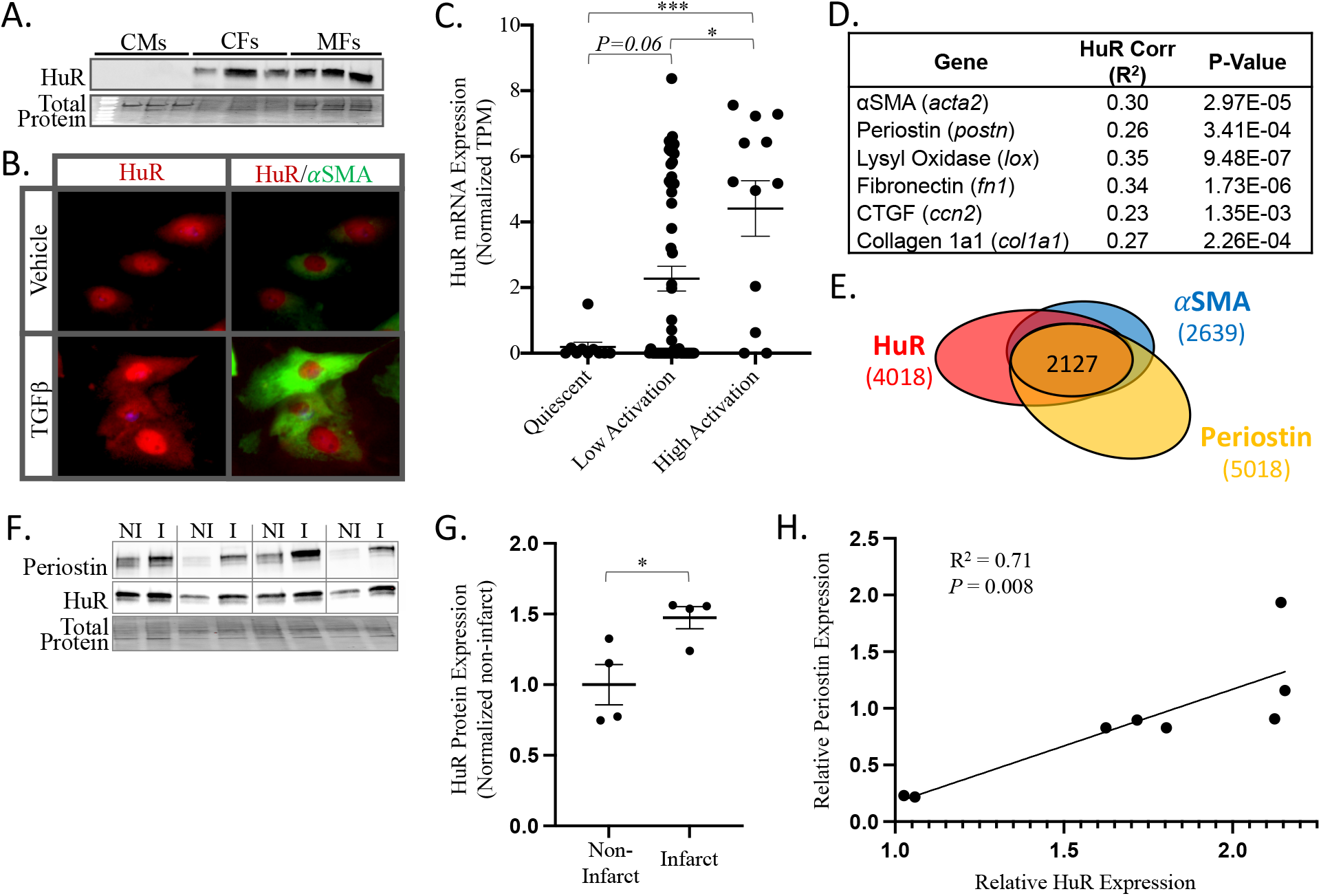
Western blotting shows that HuR expression in both CFs and MFs is significantly elevated compared to healthy cardiac myocytes (CMs) (A). Immunofluorescence shows increased HuR cytoplasmic translocation in fibroblasts at 24 hours post TGFβ treatment (B). Single-cell RNA-seq data from 185 individual murine cardiac fibroblasts isolated at 14 days post-MI shows that HuR mRNA expression is increased in fibroblasts upon activation (quantitative binning as quiescent, low, and high activation states done using gene expression levels of known myofibroblast marker genes) (C). A transcriptome wide Pearson correlation identified 4018 genes whose expression significantly correlated with HuR, of which 2127 also showed significant correlation with the MF markers *α*SMA and periostin (D). HuR shows strong expression correlation with several MF marker genes in cardiac fibroblasts (E). Western blotting for HuR on protein isolated from fibroblasts in the non-infarcted region (NI) and infarct region (I) of the mouse heart 2 weeks after 30 min ischemia followed by reperfusion supports that HuR expression increases in the fibroblasts from the infarcted region (F-G). Correlation of normalized HuR and Periostin protein expression in fibroblasts isolated from the infarct zone of post-ischemic hearts (H). \**P*≤0.05. \*\*\**P*≤0.001.

Analysis of RNA expression using a publicly available data set (Gene Expression Omnibus GSE83337) of single cell RNA seq data from 185 cardiac fibroblasts (tcf21^+^ lineage) isolated from mouse hearts at two weeks post-MI shows that HuR gene expression significantly increases upon activation to MFs (Fig. 1C). We also applied transcriptome wide correlation analysis to this data set to identify genes that correlate and co-express with HuR in cardiac fibroblasts (Fig. 1D). This was done using the weighted gene co-expression network analysis (WGCNA) R package [26,27], which has been shown as a powerful tool to predict causal relationships between gene expression networks and phenotypes [28–30]. This approach confirms significant co-expression correlation between HuR and known MF markers, including *α*SMA and periostin (Fig. 1D). In sum, we found 4018 genes that showed significant co-expression with HuR in cardiac fibroblasts, with a significant proportion of these genes (2127/4018; 53%) also showing significant co-expression with both *α*SMA and periostin (Fig. 1E) (Table S1). Taken together, this data suggests HuR activation and expression to be a strong marker of MF activity.

HuR protein expression has been previously shown to increase in the heart following myocardial infarction [31]. To determine if HuR increases specifically in fibroblasts following infarction, HuR protein expression was assessed from fibroblasts isolated from the non-infarct and infarct regions of wild-type mice at 14 days post-ischemia/reperfusion injury (Fig. 1F). Results show a significant increase in HuR expression in fibroblasts isolated from the infarct zone (Fig. 1G). Expression of periostin was also assessed as a marker of MF activity, and a significant positive correlation of HuR and periostin protein expression was observed in the isolated fibroblast fractions (Fig. 1F and 1H).

### HuR activity is necessary for TGFβ-mediated gene expression and ECM production

To identify the functional role of HuR in myofibroblast activity, primary mouse cardiac fibroblasts were treated with TGFβ (10 ng/ml for 24 hours) in the presence of KH-3 (10 *μ*M), a pharmacological inhibitor of HuR that we have previously demonstrated effective at reducing pathological cardiac remodeling in response to pressure overload [3,32]. TGFβ induced a significant increase in both periostin and collagen 1a1 expression that was completely abrogated upon HuR inhibition (Fig. 2A-B).

**Figure 2.**
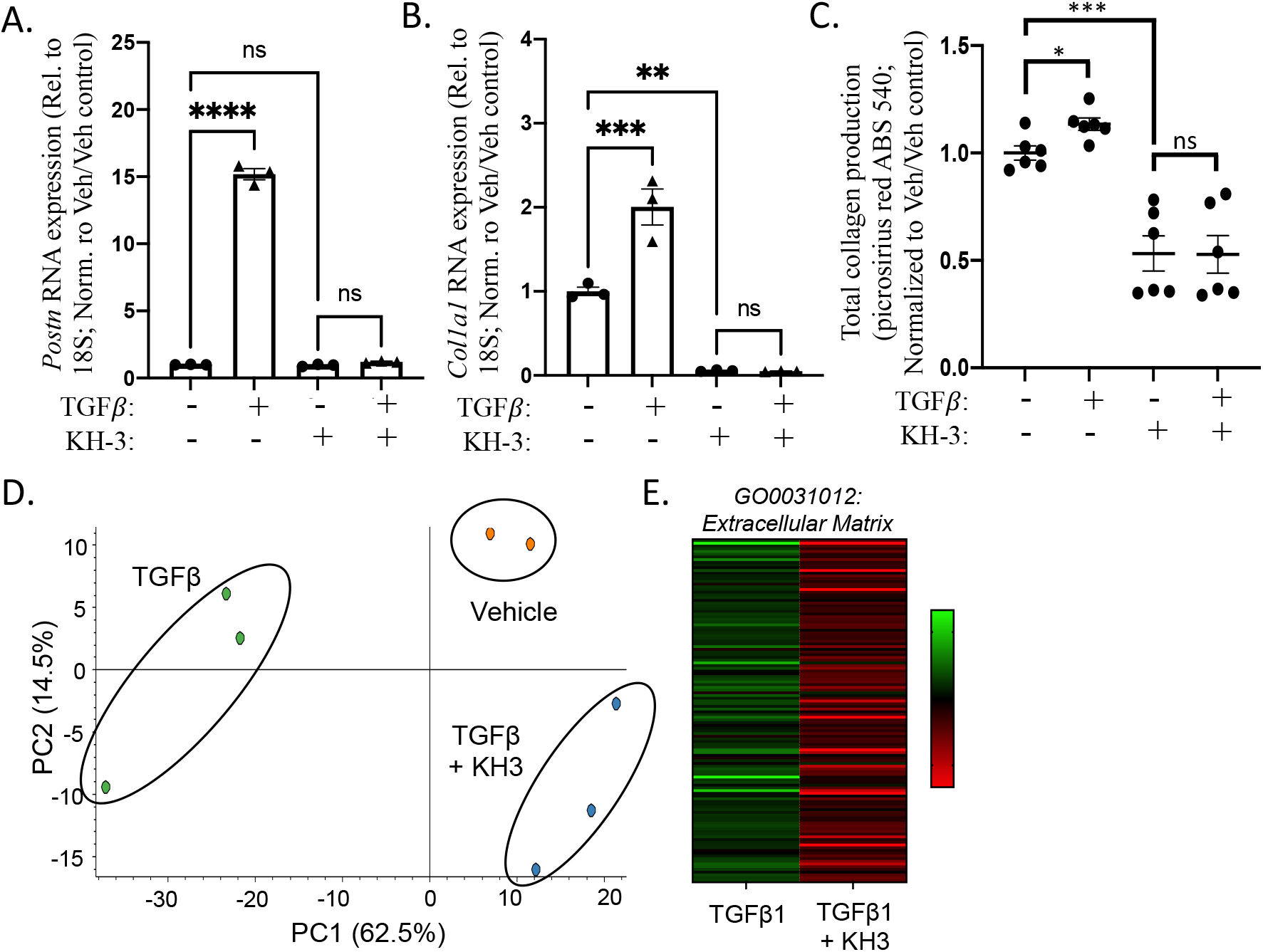
TGFβ-induced expression of periostin (*postn*) and collagen (*col1a1*) mRNA expression is blunted upon HuR inhibition via KH-3 as determined by qPCR (A-B). Primary cardiac fibroblasts treated with TGFβ, KH-3, or Vehicle for 72 hours followed by decellularization, picrosirius red staining, and quantification of total collagen show that HuR inhibition reduces total collagen production (C). Principle component analysis (PCA) plot following mass-spec analysis shows distinct grouping of Vehicle, TGFβ, or samples (D). Heat map analysis shows a large reduction in expression of ECM-related proteins following KH-3 treatment (E). \**P*≤0.05. \*\**P*≤0.01. \*\*\**P*≤0.001.

To determine if this translated to a HuR-dependent effect on total ECM production, CFs were cultured for 72 hours in the presence of TGFβ and/or KH-3 followed by decellularization and picrosirius red staining of residual ECM. Similar to gene expression, HuR inhibition significantly reduced total ECM production of cardiac fibroblasts (Fig. 2C). Interestingly, HuR inhibition also reduced the gene expression and secretion of collagen in non-stimulated CFs (Fig. 2B-C).

In addition, unbiased proteomics analysis was performed on the decellularized ECM using mass-spectrometry as previously described [33]. Principal component analysis shows a distinct sample grouping for TGFβ vs. TGFβ + KH-3 treated samples, with KH-3 treatment resulting in a proteome shift back toward that of the vehicle group (Fig. 2D). In total, 185 proteins were found to be differentially expressed within the ECM of TGFβ-treated fibroblasts in KH-3 vs. vehicle treated groups (significance defined by fold change ≥ 2 and p ≤ 0.05). Table S2). Of these 185 differentially expressed proteins, all but 2 were significantly decreased in expression following HuR inhibition, which is shown by heat map representation of significantly decreased protein expression within the *Extracellular Matrix* gene ontology group (GO0031012; Fig. 2E). These results are consistent with a decrease in ECM-related protein expression and secretion upon HuR inhibition.

### HuR inhibition blunts the myofibroblast wound healing response and contractility

Primary cardiac fibroblasts were grown in culture for 3 days without passaging prior to a scratch wound healing assay in the presence of KH-3 or vehicle control. Quantification of time lapse imaging shows that vehicle treated fibroblasts close the wound to ~50% of original scratch width (51.2 ± 16.3%) at 48 hours post-scratch, whereas fibroblasts treated with KH-3 only closed to ~90% (88.2 ± 9.7%) (Fig. 3A-B). Importantly, the blunted wound healing response is also shown upon siRNA-mediated knockdown of HuR expression, increasing confidence in the specificity of these results for HuR inhibition (Fig. 3A-B). In addition, KH-3 treated cells display decreased αSMA expression at 48 hours post-scratch compared to vehicle treated controls (Fig. 3C). Additionally, HuR inhibition significantly blunted mRNA expression of pro-fibrotic genes collagen 1a1 (*col1a1*), periostin (*postn*), fibronectin (*fn*), and TGFβ1 (*tgfb1*) at 48 hours post-scratch (Fig. 3D). We also show that KH-3 treatment did not affect the number of Hoechst positive nuclei, suggesting the HuR-dependent effect to be through modulation of cell migration, rather than proliferation (Fig. 3E). This data suggests that HuR activation is necessary to induce transition of fibroblasts into active myofibroblasts.

**Figure 3.**
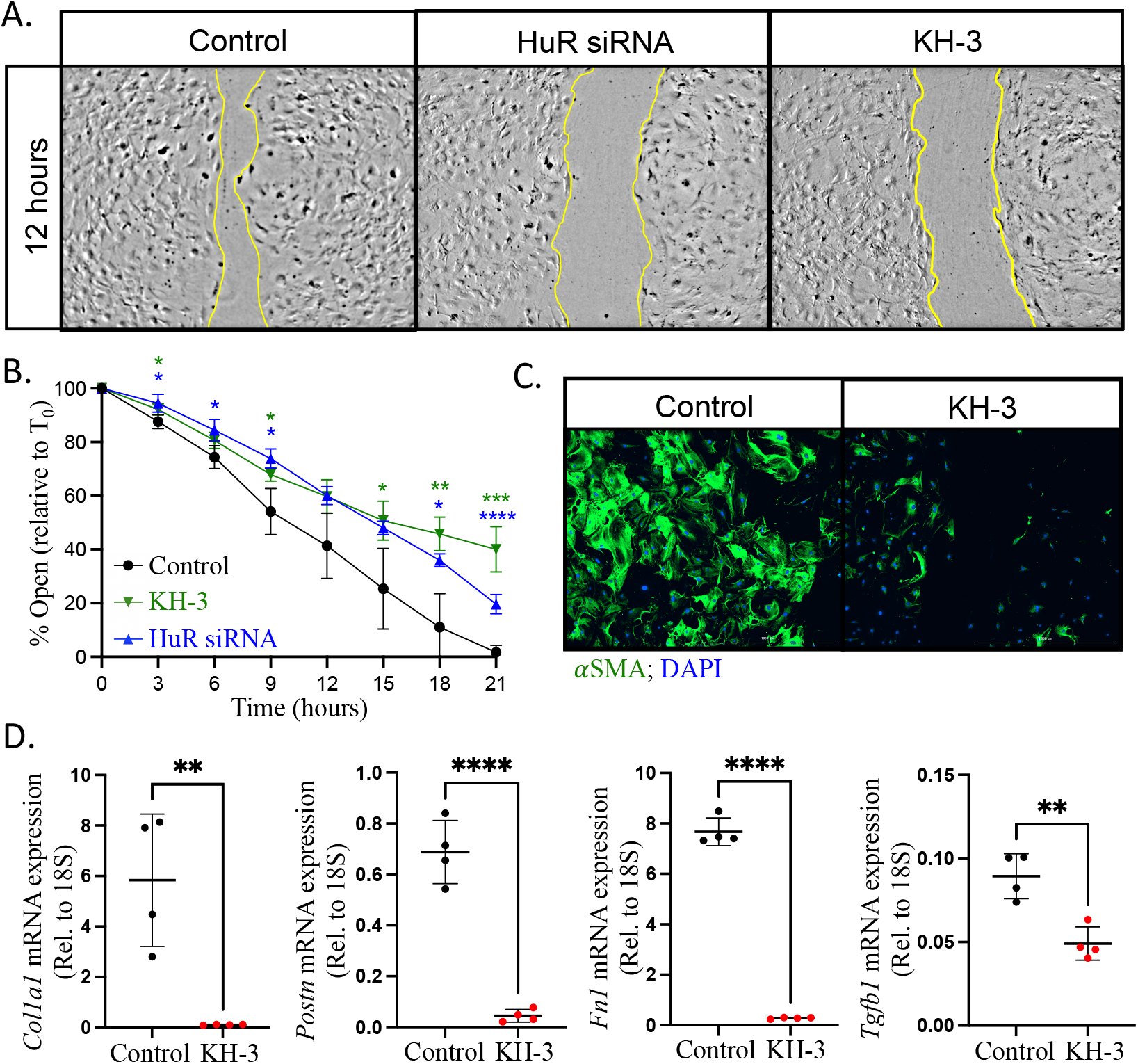
Primary adult mouse fibroblasts show a blunted wound healing response in a scratch assay upon siRNA-mediated HuR knockdown or inhibition via KH-3 (A-B). Treatment with KH-3 also inhibited the expression of induction of *α*SMA protein expression (C) as well as other pro-fibrotic ECM remodeling genes (Col1a1, Periostin, Fibronectin, and TGFβ1) at 48 hours post-scratch assay (D). \**P*≤0.05. \*\**P*≤0.01. \*\*\**P*≤0.001. \*\*\*\**P*≤0.0001.

We assessed the contractile phenotype of primary isolated fibroblasts as they transition to a myofibroblast via collagen gel contraction assay. Primary fibroblasts were seeded in free-floating collagen gels and treated for 72 hours with either TGFβ (10 ng/ml) to induce the myofibroblast contractile phenotype, KH-3 (10 *μ*M) to inhibit HuR, or both. As expected, results show that TGFβ significantly induced gel contraction (73.2 ± 9.2%) compared to vehicle only treated cells (31.7 ± 7.9%) (Fig. 4). Inhibition of HuR significantly reduced TGFβ-mediated gel contraction (38.5 ± 5.7%; *p ≤* 0.05 vs. TGFβ alone). HuR inhibition had no apparent effect on the gel contraction in unstimulated cells (41.4 ± 8.2%; not significant vs. vehicle only).

**Figure 4.**
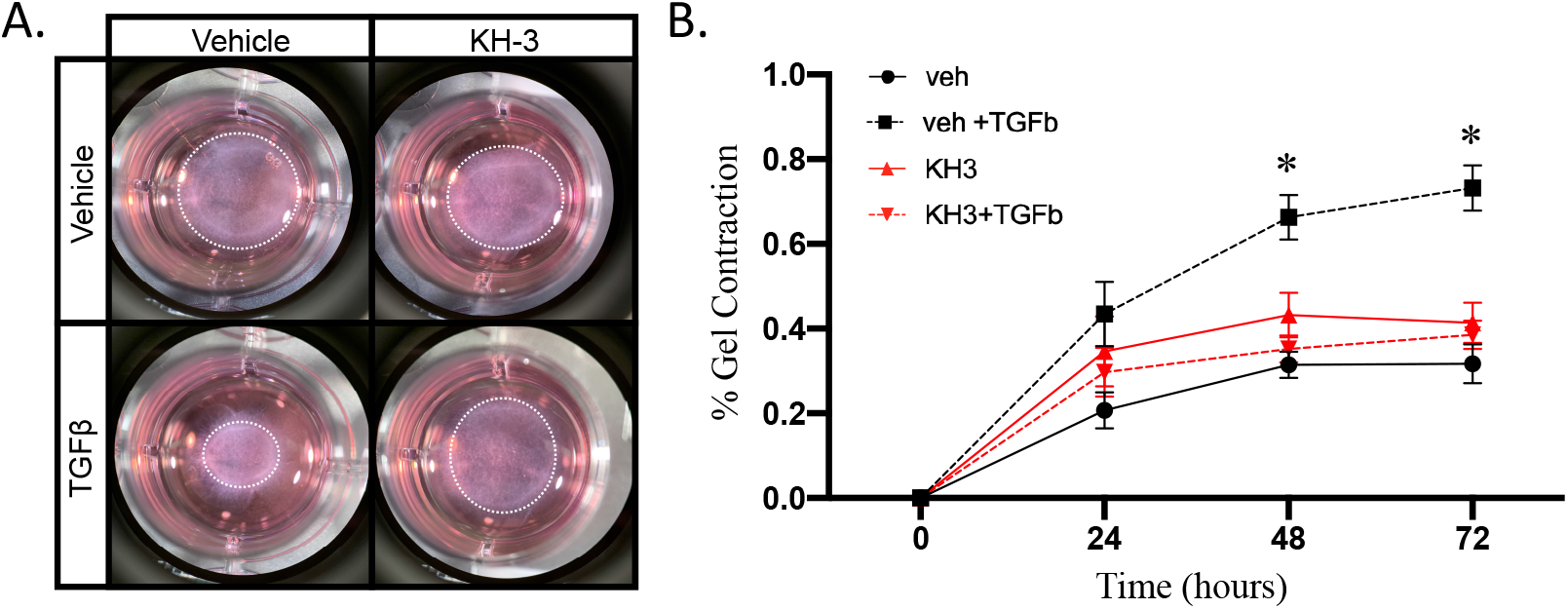
Collagen gels were seeded with cardiac fibroblasts isolated from 12 week old male mice. The gels were suspended in media with or without KH-3 and 10ng/mL TGFβ, and the rate of contraction was measured over 72 hours (A). Representative gel images at 72 hours; gels outlined by dashed line (B). N=3/group. \**P*≤0.05.

### HuR directly binds Wisp1 mRNA and regulates its expression in a TGFβ-dependent manner

To delineate HuR-dependent mechanisms of MF activity, we next used photoactivatable ribonucleoside-enhanced crosslinking and immunoprecipitation (PAR-CLIP) to identify directly bound HuR target genes in primary cardiac fibroblasts. Following PAR-CLIP, total HuR bound RNA was isolated and identified via RNA-sequencing. Results identified 1821 HuR-bound transcripts (Table S3) in quiescent CFs, and 24 transcripts that showed greater than a two-fold increase in HuR binding following TGFβ treatment for 24 hours (Fig. 5A; Table S4). Of these 24 TGFβ-enriched HuR targets, 11 of these genes are also significantly co-expressed with HuR, periostin, and *α*SMA gene expression in cardiac fibroblasts (Fig. 5A red circle and Fig 5B table; co-expression correlations were derived from the scRNA-seq data set used in Fig. 1). Wisp1 was identified as the TGFβ-enriched HuR target gene that most strongly correlated with HuR co-expression in MFs (Fig. 5B). Confirmation of direct HuR binding to Wisp1 mRNA was done via qPCR and shows a low level of binding in unstimulated CFs that is significantly increased following TGFβ treatment (Fig. 5C). To confirm a functional effect for HuR binding on Wisp1 expression, results show that the TGFβ-mediated increase in Wisp1 mRNA expression is completely abrogated by HuR inhibition (Fig. 5D).

**Figure 5.**
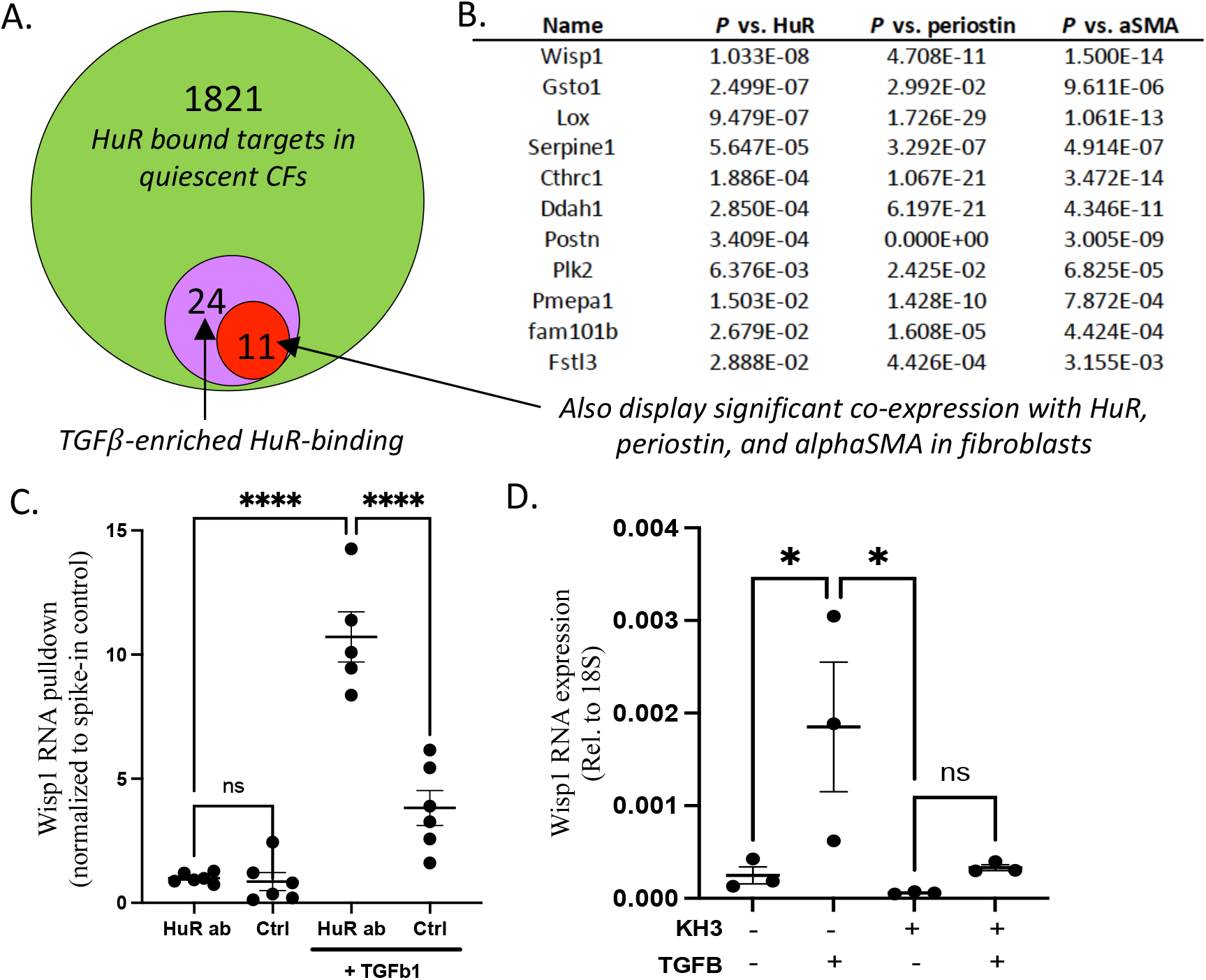
PAR-CLIP identified 1822 HuR bound target genes in CFs, 24 of which showed increased HuR binding at 24 hours post TGFβ treatment (**A**). Of these 24 genes, 11 showed significant co-expression with HuR, periostin, and *α*SMA (**B**; correlation p-values displayed). Direct HuR binding to Wisp1 in primary CFs was confirmed via qPCR validation of PAR-CLIP results using HuR or control antibody and TGFβ (10ng/ml for 24 hours) or vehicle control (**C**). HuR inhibition via KH-3 reduces TGFβ-dependent (10ng/ml for 24 hours) expression of Wisp1 in primary CFs (**D**).

### HuR-dependent Wisp1 expression mediates MF functional activity

To determine the functional role of Wisp1 expression downstream of HuR, recombinant Wisp1 was added to primary CFs in a collagen gel contraction assay concomitant with KH-3-mediated HuR inhibition. Results show that Wisp1 is able to partially rescue the HuR-dependent contractile function of MFs (Fig. 6A). Results also show that HuR inhibition via KH-3 has no inhibitory effect on gel contraction induced by recombinant Wisp1 alone (Fig. 6B). Together, these data demonstrate a functional role for HuR-dependent Wisp1 expression in TGFβ-induced MF activity and identify Wisp1 as an independent downstream mediator of HuR regulation in MFs (Fig. 7).

**Figure 6.**
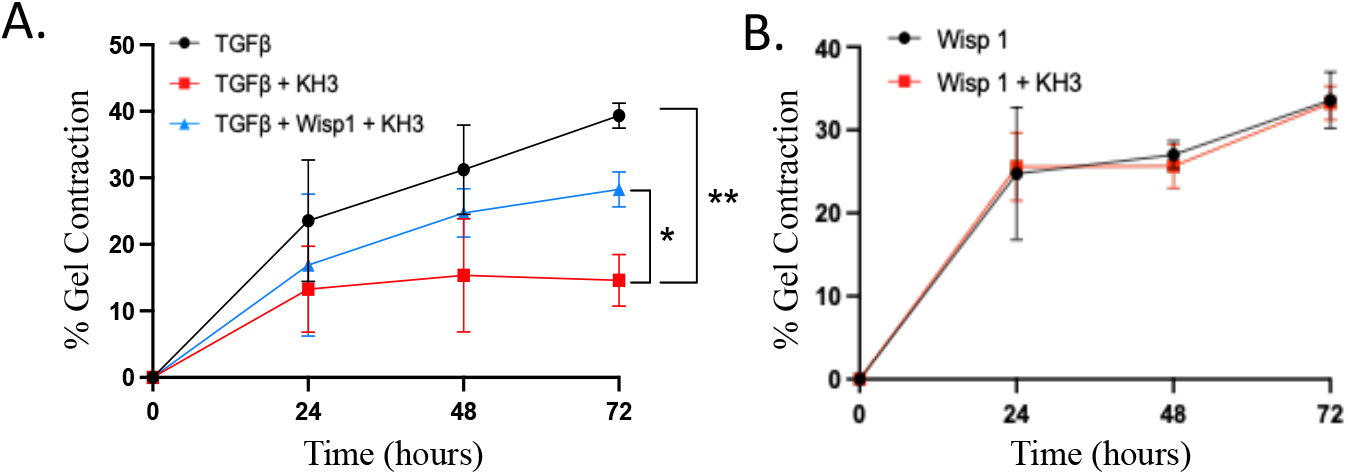
Recombinant Wisp1 is able to rescue the KH-3-meidated suppression of TGFβ-mediated gel contraction in primary CFs (**A**), but HuR inhibition itself has no impact on Wisp1-mediated gel contraction (**B**). \**P*≤0.05. \*\**P*≤0.01. \*\*\**P*≤0.0001. N ≥ 3/group.

**Figure 7.**
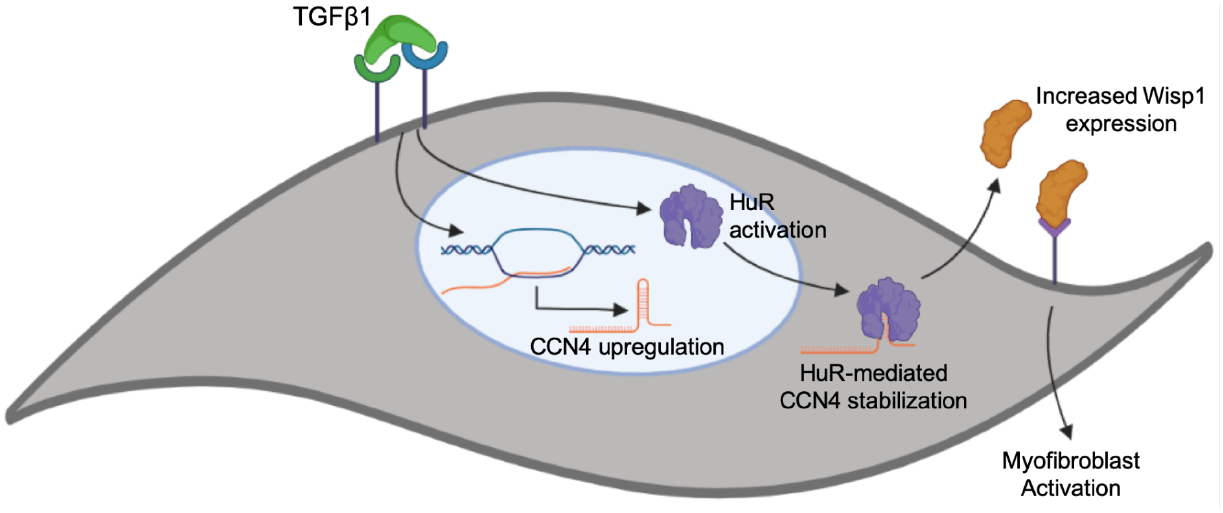
Working model of myofibroblast activation via HuR-dependent expression of Wisp1 downstream of TGFβ stimulation.

## DISCUSSION

This is the first work to show that HuR plays a direct functional role in TGFβ-mediated activation of cardiac fibroblasts. We have previously shown that pharmacological inhibition of HuR reduces pathological cardiac remodeling in response to pressure overload, including a specific role for HuR in cardiac myocytes [3,9]. However, in this work, we show a high level of HuR mRNA and protein expression in isolated CFs and MFs, strongly suggesting a functional role in the ECM remodeling activity of these cells. Bai et al previously showed TGFβ-mediated HuR activation in rat cardiac fibroblasts that was necessary for TGFβ-dependent increases in *tgfb1*, *col1a1*, *col1a3*, and *fn* expression, but did not directly assess the functional consequences of this HuR-dependent gene expression [8]. We have utilized functional assays to delineate the HuR-dependent effects of fibroblast migration (scratch assay), ECM remodeling (proteomics), and contractile function (collagen gel contraction) to conclusively show a necessary role for HuR in MF activity in response to TGFβ. Furthermore, we employed a bioinformatics approach using both scRNA-seq data and PAR-CLIP to identify directly bound HuR target genes and have identified Wisp1 as a functional downstream effector of HuR in MFs.

In this work, we focused primarily on Wisp1 as a HuR target gene whose binding was significantly enriched in a TGFβ-dependent manner, and went on to show a functional role for Wisp1 expression in MF activity downstream of HuR. However, our RNA sequencing results of HuR targets suggests direct HuR binding to additional genes commonly associated with the MF phenotype, such as *lox* and *postn* (Fig. 5B). Our results clearly demonstrate a critical dependency for HuR activity for CF activation to a MF phenotype at the level of both function and gene expression. However, it remains unclear how broadly HuR orchestrates expression of additional MF marker genes or those needed for cell activation/differentiation. Interestingly, if we examine function of HuR bound target genes in CFs by gene ontology (GO) analysis, results show an enrichment for GO groups associated with mRNA processing, translation, and Golgi-mediated protein secretion (Table S5). These processes would obviously be critical to CF activation and the ECM remodeling functions of MFs, but also suggest a broader role for HuR in the ability of CFs to respond to a pro-fibrotic stimulus. This is in agreement with existing work showing a critical role for HuR activation in response to stress and growth stimuli in other cell types, including the established function of HuR in mediating tumorigenesis [34–36]. Additionally, our previously published work showed no effect of HuR deletion in quiescent, terminally differentiated adult cardiac myocytes [3], but HuR knockdown potently reduced hypertrophic growth in neonatal myocytes [9]. This pleiotropic role for HuR and the breadth of mRNA targets across cell types and physiological conditions emphasizes the importance of identifying specific HuR bound targets in pathogenic conditions [37,38]. For example, our results identified 1821 HuR bound targets in CFs, but only a small number of these (24, including Wisp1) showed enriched binding in response to TGFβ, suggesting this pool of genes as the specific downstream mediators of HuR function in MF activation.

WNT1-inducible signaling pathway protein 1 (Wisp1/*Ccn4*) belongs to the connective tissue growth factor (CTGF) and nephroblastoma overexpressed (CCN) family of proteins that have been shown to have potent effects on differentiation, proliferation, and ECM remodeling in multiple cell types [39]. Wisp1 expression has been shown to be increased in tumor cells as well as under stress or injury conditions such as ischemic brain injury, spinal cord injury, or following oxidative stress [40,41]. Wnt and TGFβ signaling interact in pro-fibrotic pathways of fibroblast activation, and Wisp1 expression has been previously shown to be modulated by both TGFβ and Wnt [42,43]. Inhibition of Wnt signaling has been shown using both *in vitro* and *in vivo* models to reduce TGFβ-mediated fibroblast activation [44,45]. Specifically, Wisp1 expression has been shown to increase in the myocardium post-MI, and has been shown using *in vitro* models to independently promote both hypertrophy and survival in myocytes as well as collagen production in fibroblasts [11,46]. Wisp1 expression has also been suggested to be a strong marker of active MFs [25], and our results here demonstrate Wisp1 to be a functional HuR-dependent mediator of MF activity.

Interestingly, we also observed a HuR-dependent increase in total cardiac Wisp1 mRNA expression following pressure overload in our previous work using myocyte-specific HuR^-/-^ mice [3]. However, it’s unclear to what extent the reduced pathology from myocyte-specific deletion of HuR or pan-inhibition across myocardial cell types using KH-3 may be a result of reductions in HuR-dependent Wisp1 expression. As a secreted protein that has been shown to have independent effects on myocytes and fibroblasts, the primary source of functional Wisp1 expression in the myocardium is also unclear, though secretion by active MFs would appear most likely. Indeed, we did observe a reduction in fibrosis and numerous MF markers in both pharmacological inhibition and myocyte-specific HuR^-/-^ mice, suggestive of reduced MF activity in both models. Unfortunately, the functional role of Wisp1 *in vivo* has not been well explored, in part due to the lack of genetic models allowing for cell-type specific deletion. Mice with a whole-body deletion of Wisp1 were shown to have decreased bone mineral density, which was suggested to be due to a decrease in bone collagen deposition, but the impact of Wisp1 deletion on cardiac physiology remains unknown [41].

In summary, we demonstrate that HuR-dependent expression of Wisp1 is critical for TGFβ-dependent activation of cardiac fibroblasts to a myofibroblast phenotype (Fig. 7). The fibroblast specific function of HuR-Wisp1 signaling using *in vivo* models is needed to further elucidate the functional mechanisms of this pathway in cardiac remodeling. We have previously demonstrated that pharmacological inhibition of HuR ameliorates pathological cardiac remodeling following pressure overload in mice, and these results suggest HuR-dependent MF activation as a potential mechanism and suggest Wisp1 may represent a more specific downstream target to reduce MF activity.

## DATA AVAILABILITY STATEMENT

The single-cell RNA-seq dataset used in this work has been previously published and is publicly available in the NIH Gene Expression Omnibus as project #GSE83337[25]. Full sequencing results for HuR bound targets via PAR-CLIP are available in the NIH Gene Expression Omnibus (GSE193931).

## REFERENCES

[1] R. Bretherton, D. Bugg, E. Olszewski, J. Davis, Regulators of cardiac fibroblast cell state, Matrix Biol. 91-92 (2020) 117–135. doi:10.1016/j.matbio.2020.04.002.

[2] J.G. Travers, F.A. Kamal, J. Robbins, K.E. Yutzey, B.C. Blaxall, Cardiac Fibrosis: The Fibroblast Awakens, Circ. Res. 118 (2016) 1021–1040. doi:10.1161/CIRCRESAHA.115.306565.

[3] L.C. Green, S.R. Anthony, S. Slone, L. Lanzillotta, M.L. Nieman, X. Wu, et al., Human antigen R as a therapeutic target in pathological cardiac hypertrophy, JCI Insight. 4 (2019) 47. doi:10.1172/jci.insight.121541.

[4] H. Khalil, O. Kanisicak, V. Prasad, R.N. Correll, X. Fu, T. Schips, et al., Fibroblast-specific TGF-β-Smad2/3 signaling underlies cardiac fibrosis, J. Clin. Invest. 127 (2017) 3770–3783. doi:10.1172/JCI94753.

[5] C. Humeres, N.G. Frangogiannis, Fibroblasts in the Infarcted, Remodeling, and Failing Heart, JACC Basic Transl Sci. 4 (2019) 449–467. doi:10.1016/j.jacbts.2019.02.006.

[6] A. Hanna, N.G. Frangogiannis, The Role of the TGF-β Superfamily in Myocardial Infarction, Front Cardiovasc Med. 6 (2019) 140. doi:10.3389/fcvm.2019.00140.

[7] R.M. Burke, K.N. Burgos Villar, E.M. Small, Fibroblast contributions to ischemic cardiac remodeling, Cell. Signal. 77 (2021) 109824. doi:10.1016/j.cellsig.2020.109824.

[8] D. Bai, Q. Gao, C. Li, L. Ge, Y. Gao, H. Wang, A conserved TGFβ1/HuR feedback circuit regulates the fibrogenic response in fibroblasts, Cell. Signal. 24 (2012) 1426–1432. doi:10.1016/j.cellsig.2012.03.003.

[9] S. Slone, S.R. Anthony, X. Wu, J.B. Benoit, J. Aubé, L. Xu, et al., Activation of HuR downstream of p38 MAPK promotes cardiomyocyte hypertrophy, Cell. Signal. 28 (2016) 1735–1741. doi:10.1016/j.cellsig.2016.08.005.

[10] P.K. Govindappa, M. Patil, V.N.S. Garikipati, S.K. Verma, S. Saheera, G. Narasimhan, et al., Targeting exosome-associated human antigen R attenuates fibrosis and inflammation in diabetic heart, The FASEB Journal. 34 (2020) 2238–2251. doi:10.1096/fj.201901995R.

[11] J.T. Colston, S.D. de la Rosa, M. Koehler, K. Gonzales, R. Mestril, G.L. Freeman, et al., Wnt-induced secreted protein-1 is a prohypertrophic and profibrotic growth factor, Am. J. Physiol. Heart Circ. Physiol. 293 (2007) H1839–46. doi:10.1152/ajpheart.00428.2007.

[12] B. Venkatesan, S.D. Prabhu, K. Venkatachalam, S. Mummidi, A.J. Valente, R.A. Clark, et al., WNT1-inducible signaling pathway protein-1 activates diverse cell survival pathways and blocks doxorubicin-induced cardiomyocyte death, Cell. Signal. 22 (2010) 809–820. doi:10.1016/j.cellsig.2010.01.005.

[13] D. Smiley, M.A. Smith, V. Carreira, M. Jiang, S.E. Koch, M. Kelley, et al., Increased fibrosis and progression to heart failure in MRL mice following ischemia/reperfusion injury, Cardiovasc. Pathol. 23 (2014) 327–334. doi:10.1016/j.carpath.2014.06.001.

[14] J.G. Travers, F.A. Kamal, I. Valiente-Alandi, M.L. Nieman, M.A. Sargent, J.N. Lorenz, et al., Pharmacological and Activated Fibroblast Targeting of Gβγ-GRK2 After Myocardial Ischemia Attenuates Heart Failure Progression, J. Am. Coll. Cardiol. 70 (2017) 958–971. doi:10.1016/j.jacc.2017.06.049.

[15] A.R. Pinto, A. Ilinykh, M.J. Ivey, J.T. Kuwabara, M.L. D’Antoni, R. Debuque, et al., Revisiting Cardiac Cellular Composition, Circ. Res. 118 (2016) 400–409. doi:10.1161/CIRCRESAHA.115.307778.

[16] S.R. Anthony, A. Guarnieri, L. Lanzillotta, A. Gozdiff, L.C. Green, K. O’Grady, et al., HuR expression in adipose tissue mediates energy expenditure and acute thermogenesis independent of UCP1 expression, Adipocyte. 9 (2020) 335–345. doi:10.1080/21623945.2020.1782021.

[17] B. Rivero-Gutiérrez, A. Anzola, O. Martínez-Augustin, F.S. de Medina, Stain-free detection as loading control alternative to Ponceau and housekeeping protein immunodetection in Western blotting, Anal Biochem. 467 (2014) 1–3. doi:10.1016/j.ab.2014.08.027.

[18] K.J. Livak, T.D. Schmittgen, Analysis of relative gene expression data using real-time quantitative PCR and the 2(-Delta Delta C(T)) Method, Methods. 25 (2001) 402–408. doi:10.1006/meth.2001.1262.

[19] A. Suarez-Arnedo, F. Torres Figueroa, C. Clavijo, P. Arbeláez, J.C. Cruz, C. Muñoz-Camargo, An image J plugin for the high throughput image analysis of in vitro scratch wound healing assays, PLoS ONE. 15 (2020) e0232565. doi:10.1371/journal.pone.0232565.

[20] N. Pincha, D. Saha, T. Bhatt, R.K. Zirmire, C. Jamora, Activation of Fibroblast Contractility via Cell-Cell Interactions and Soluble Signals, Bio Protoc. 8 (2018) e3021–e3021. doi:10.21769/BioProtoc.3021.

[21] P. Vavken, S. Joshi, M.M. Murray, TRITON-X is most effective among three decellularization agents for ACL tissue engineering, J Orthop Res. 27 (2009) 1612–1618. doi:10.1002/jor.20932.

[22] Q. Xu, J.T. Norman, S. Shrivastav, J. Lucio-Cazana, J.B. Kopp, In vitro models of TGF-beta-induced fibrosis suitable for high-throughput screening of antifibrotic agents, Am J Physiol Renal Physiol. 293 (2007) F631–40. doi:10.1152/ajprenal.00379.2006.

[23] M. Fisher, R.A. Jones, L. Huang, J.L. Haylor, M. El Nahas, M. Griffin, et al., Modulation of tissue transglutaminase in tubular epithelial cells alters extracellular matrix levels: a potential mechanism of tissue scarring, Matrix Biol. 28 (2009) 20–31. doi:10.1016/j.matbio.2008.10.003.

[24] A.R. Guarnieri, S.R. Anthony, A. Gozdiff, L.C. Green, S.M. Fleifil, S. Slone, et al., Adipocyte-specific deletion of HuR induces spontaneous cardiac hypertrophy and fibrosis, Am. J. Physiol. Heart Circ. Physiol. 321 (2021) H228–H241. doi:10.1152/ajpheart.00957.2020.

[25] O. Kanisicak, H. Khalil, M.J. Ivey, J. Karch, B.D. Maliken, R.N. Correll, et al., Genetic lineage tracing defines myofibroblast origin and function in the injured heart, Nat Commun. 7 (2016) 12260–14. doi:10.1038/ncomms12260.

[26] P. Langfelder, S. Horvath, WGCNA: an R package for weighted correlation network analysis, BMC Bioinformatics. 9 (2008) 559–13. doi:10.1186/1471-2105-9-559.

[27] B. Zhang, S. Horvath, A general framework for weighted gene co-expression network analysis, Stat Appl Genet Mol Biol. 4 (2005) Article17. doi:10.2202/1544-6115.1128.

[28] B.E. Haas, S. Horvath, K.H. Pietiläinen, R.M. Cantor, E. Nikkola, D. Weissglas-Volkov, et al., Adipose co-expression networks across Finns and Mexicans identify novel triglyceride-associated genes, BMC Med Genomics. 5 (2012) 61–10. doi:10.1186/1755-8794-5-61.

[29] X. Wang, J. Duanmu, X. Fu, T. Li, Q. Jiang, Analyzing and validating the prognostic value and mechanism of colon cancer immune microenvironment, J Transl Med. 18 (2020) 324–14. doi:10.1186/s12967-020-02491-w.

[30] M. Wang, J. Wang, J. Liu, L. Zhu, H. Ma, J. Zou, et al., Systematic prediction of key genes for ovarian cancer by co-expression network analysis, J. Cell. Mol. Med. 24 (2020) 6298–6307. doi:10.1111/jcmm.15271.

[31] P. Krishnamurthy, J. Rajasingh, E. Lambers, G. Qin, D.W. Losordo, R. Kishore, IL-10 Inhibits Inflammation and Attenuates Left Ventricular Remodeling After Myocardial Infarction via Activation of STAT3 and Suppression of HuR, Circ. Res. 104 (2009) e9–e18. doi:10.1161/CIRCRESAHA.108.188243.

[32] X. Wu, L. Lan, D.M. Wilson, R.T. Marquez, W.-C. Tsao, P. Gao, et al., Identification and Validation of Novel Small Molecule Disruptors of HuR-mRNA Interaction, ACS Chem. Biol. (2015) 150317154550000–9. doi:10.1021/cb500851u.

[33] J. Barallobre-Barreiro, A. Didangelos, F.A. Schoendube, I. Drozdov, X. Yin, M. Fernández-Caggiano, et al., Proteomics analysis of cardiac extracellular matrix remodeling in a porcine model of ischemia/reperfusion injury, Circulation. 125 (2012) 789–802. doi:10.1161/CIRCULATIONAHA.111.056952.

[34] S. Srikantan, M. Gorospe, HuR function in disease, Front Biosci (Landmark Ed). 17 (2012) 189–205. doi:10.2741/3921.

[35] J. Wang, Y. Guo, H. Chu, Y. Guan, J. Bi, B. Wang, Multiple functions of the RNA-binding protein HuR in cancer progression, treatment responses and prognosis, Int J Mol Sci. 14 (2013) 10015–10041. doi:10.3390/ijms140510015.

[36] C.W. Schultz, R. Preet, T. Dhir, D.A. Dixon, J.R. Brody, Understanding and targeting the disease-related RNA binding protein human antigen R (HuR), Wiley Interdiscip Rev RNA. 11 (2020) e1581. doi:10.1002/wrna.1581.

[37] P.J. Uren, S.C. Burns, J. Ruan, K.K. Singh, A.D. Smith, L.O.F. Penalva, Genomic analyses of the RNA-binding protein Hu antigen R (HuR) identify a complex network of target genes and novel characteristics of its binding sites, J. Biol. Chem. 286 (2011) 37063–37066. doi:10.1074/jbc.C111.266882.

[38] C. Barreau, L. Paillard, H.B. Osborne, AU-rich elements and associated factors: are there unifying principles? Nucleic Acids Res. 33 (2005) 7138–7150. doi:10.1093/nar/gki1012.

[39] K. Maiese, WISP1: Clinical insights for a proliferative and restorative member of the CCN family, Curr Neurovasc Res. 11 (2014) 378–389. doi:10.2174/1567202611666140912115107.

[40] I. Gurbuz, R. Chiquet-Ehrismann, CCN4/WISP1 (WNT1 inducible signaling pathway protein 1): a focus on its role in cancer, Int J Biochem Cell Biol. 62 (2015) 142–146. doi:10.1016/j.biocel.2015.03.007.

[41] A. Maeda, M. Ono, K. Holmbeck, L. Li, T.M. Kilts, V. Kram, et al., WNT1-induced Secreted Protein-1 (WISP1), a Novel Regulator of Bone Turnover and Wnt Signaling, J. Biol. Chem. 290 (2015) 14004–14018. doi:10.1074/jbc.M114.628818.

[42] M.H. van den Bosch, T.A. Gleissl, A.B. Blom, W.B. van den Berg, P.L. van Lent, P.M. van der Kraan, Wnts talking with the TGF-β superfamily: WISPers about modulation of osteoarthritis, Rheumatology (Oxford). 55 (2016) 1536–1547. doi:10.1093/rheumatology/kev402.

[43] S. Klee, M. Lehmann, D.E. Wagner, H.A. Baarsma, M. Königshoff, WISP1 mediates IL-6-dependent proliferation in primary human lung fibroblasts, Sci. Rep. 6 (2016) 20547–11.doi:10.1038/srep20547.

[44] F.-L. Xiang, M. Fang, K.E. Yutzey, Loss of β-catenin in resident cardiac fibroblasts attenuates fibrosis induced by pressure overload in mice, Nat Commun. 8 (2017) 712–12. doi:10.1038/s41467-017-00840-w.

[45] A. Akhmetshina, K. Palumbo, C. Dees, C. Bergmann, P. Venalis, P. Zerr, et al., Activation of canonical Wnt signalling is required for TGF-β-mediated fibrosis, Nat Commun. 3 (2012) 735–12. doi:10.1038/ncomms1734.

[46] K. Venkatachalam, B. Venkatesan, A.J. Valente, P.C. Melby, S. Nandish, J.E.B. Reusch, et al., WISP1, a pro-mitogenic, pro-survival factor, mediates tumor necrosis factor-alpha (TNF-alpha)-stimulated cardiac fibroblast proliferation but inhibits TNF-alpha-induced cardiomyocyte death, J. Biol. Chem. 284 (2009) 14414–14427. doi:10.1074/jbc.M809757200.

