## Supplementary figures and images for "HuR-dependent expression of Wisp1 is necessary for TGF*β*-induced cardiac myofibroblast activity"

### Figure S1

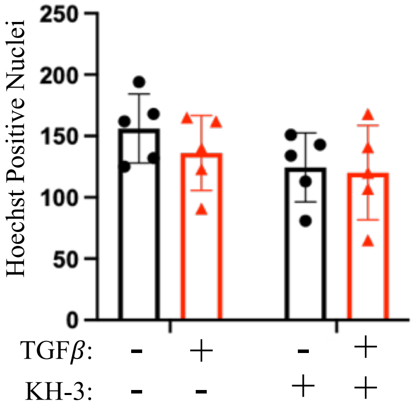
